# A spatially explicit model of pollinator-plant-pathogen interactions

**DOI:** 10.1101/2025.03.11.642689

**Authors:** Sean A. Rands, Cristina Botías, Elizabeth Nicholls, Natalie Hempel de Ibarra

## Abstract

Flowers have long been suggested to be infection hotspots for pollinator diseases, as, during their brief lifespan, they are visited and handled by many individuals from a diverse range of species. Evidence that floral hotspots are an important point of contagion is building, but few models currently allow us to characterise these novel plant-pathogen-pollinator systems where horizontal infections occur from concentrated, short-lived sites of contagion. Infection success is going to depend heavily upon the behaviour of individual pollinators, alongside the structure of the floral landscape that they are foraging within. Here, we describe an individual-based model that considers the impact of environmental heterogeneity and individual differences in movement on the spread of a disease that is spread *via* floral hotspots, considering a centrally nesting social pollinator (such as a social bee). The biggest effects we saw were associated with the likelihoods of leaving and staying in the nest and the distances travelled by the pollinators, and so it is likely that social pollinators, most of which are constrained to return to a central nest, could be strongly impacted by the schedules they use for allocating foraging behaviour. We also considered the impact of the pollinators being able to deposit and detect temporary scent-marks, which could inform an uninfected pollinator of a potentially infected site. Scent marks have a negative impact on pathogen transmission, and their effect may be dependent on both the longevity of the scent mark, the turnover of flowers in the environment, and the factors affecting pathogen viability and infectivity. Overall, although the structure of the landscape may have limited impact on pathogen spread, the behaviour of the pollinators is important, and needs further consideration within models of this pathogen system.

**AUTHOR SUMMARY:** Flowers have long been suggested to be infection hotspots for pollinator diseases, as, during their brief lifespan, they are visited and handled by many individuals from a diverse range of species. Evidence that floral hotspots are an important point of contagion is building, but we currently know little about the environmental biology of these infections. We describe a model that considers how pathogens caught at floral infection hotspots might spread in populations of social pollinators (such as a bumblebee or honeybee), and show that both pollinator behaviour and how flowers are distributed through the landscape can have a big impact on the spread of the pathogen. Pollinators can leave temporary detectable scent marks on the surfaces of the flowers that they visit: our model demonstrates that being able to detect these marks reduces the spread of the pathogen, as the pollinator is better able to avoid potential hotspots.

## INTRODUCTION

Many pathogens rely on hotspots of contagion to successfully spread between individuals, where the pathogen sits in the environment and waits for a potential host to come within a suitable range for the pathogen to infect it. For example, this is seen in horizontally-transmitted pathogens that rely on faecal-oral contagion [1], droplet contamination of surfaces [2], and also in the strategies of small mobile parasitoids and parasites with limited range such as entomopathogenic nematodes and mites [3]. Contagion will be enhanced if the pathogen is located at the specific parts of the surface that an animal is more likely to make contact with. In the case of flowers, this could be around the site of the reward (such as nectar) provided to visiting pollinators. This means that plant-pollinator interactions present an excellent opportunity for studying how spatially-concentrated hotspots of contagion can impact on their hosts [4,5]. Co-evolution has shaped tightly-orchestrated visits to flowers, where the pollinator often has to approach, orientate and interact with particular structures in order to access and extract a reward, which may in turn require the pollinator to remain in contact with specific areas of the floral substrate for extended periods of time. Over the ‘lifetime’ of an individual flower, multiple visitors may place their mouthparts in the same reward-providing region of the flower, or sit, groom or defecate [6,7] on the surface of the flower. In many species, evolutionary processes have shaped the morphology of flowers to be easily detectable and accessible, which encourages multiple visitors, which in turn may be associated with enhancing pathogen transmission [5,8–11]. However, we still know relatively little about these relationships, and only recently a couple of models have addressed this pathway fitted around measured transmission effects [12–14]. These previous models implement links between pollinator networks and infection considering visitation frequencies of pollinators. In this paper, we present a complementary approach with a model that explicitly incorporates temporal and spatial parameters derived from behavioural and sensory principles of foraging pollinators. As pollinator behaviour is highly dependent upon the spatial configuration of the pollinator’s landscape, we assume that these behavioural parameters will drive infection dynamics that can be captured with an Individual-based modelling approach, bringing movement rules, landscape form and population processes together.

Here, we focus on flower-pollinator interactions involving bees. Bees are a significant and abundant group of pollinators, and the foraging and navigational behaviours of bees have been studied for a long time [15–20]. Flowers are known to act as points of environmental transmission for a range of different bee-infecting pathogens. Trypanosomes *Crithidia bombi* are spread between bumblebees *Bombus* spp. *via* the faecal-oral route on floral surfaces [4,21–23], and microsporidia (*Nosema bombi*) and neogregarines (*Apicystic bombi*) can also be deposited and then vectored *via* flowers [23]. Mites can use flowers as staging posts to transfer between bumblebees [24], and evidence suggests that this may be a route for *Varroa* transmission [25]. Some RNA viruses can be deposited on flowers by visiting honeybees *Apis mellifera* [26], and these deposited viruses are capable of infecting other hymenopteran pollinators [27–29]. Some fungal diseases may also be transmitted through flower-sharing [30]. It is also worth noting that pollinators may be vectors for organisms that affect plant health, both positively [31] and negatively [32].

Floral contagion hotspots offer an interesting infection system which is ecologically very different to many other environmental contagion systems that have been studied. Floral visitors are obliged to visit flowers in order to harvest rewards, which is arguably similar to obligate contagion sites like watering holes. However, unlike other obligate sites, there is likely to be a very large amount of choice with multiple flowers of different species being present. Flowers are also transient, and an individual contaminated flower may only be a viable foraging site for a matter of hours or days, and may only be available for intermittent short periods of the day if it closes at night. Furthermore, similar to animals visiting other obligate contagion sites, visitors may have behavioural mechanisms that allow them to detect and respond adaptively to a flower that has recently been visited (and therefore potentially contaminated), such as the scent mark deposition and detection seen in bee species [5,33,34]. The pathogens at floral hotspots will therefore be subject to both intense competition and a strict time-limit on their longevity and availability. This suggests that they will either have to be distributed widely over multiple sites, or else be extremely efficient in their transmission and very resistant to environmental conditions. So far, models of pathogens transmitted *via* insect-pollinated flowers have used an ecological trait-based approach to demonstrate that floral hotspot transmission can be an effective route for pathogens. Truitt *et al.* [35] developed a compartmental model using a susceptible-infectious-susceptible framework and a related continuous trait distribution model, and explored the dynamics of a multi-species infection (where the infection can be passed between different pollinator species) that could be transmitted *via* contaminated flowers. This model principally considered how disease success may rely on multi-species interactions. Ellner *et al.* [14] expanded on this model, looking at how specialisation contributes to disease dynamics, and Burnham *et al.* [28] built on [35], and considered how different management strategies could reduce the spread of infections.

Compartmental models such as those described above reveal much about pathogen transmission dynamics, but largely fail to account for the spatial constraints imposed by pollinator biology. As such, it is not clear how floral transmission type selects for pathogen traits. In particular, the longevity of pathogens is a problem, as pollinator visits occur stochastically while pathogens on flowers are poorly protected from environmental risks. Many pollinators (including most bee species) are central-place foragers, and need to return to a central nest site to store the resources they collect and provision their young [5]. This means that they are constrained by how far they travel through the environment, which can impact on their likelihood of interacting with other individuals and species, and thus affect the probability of transmission within a critical time window that allows the pathogen to replicate and spread [5]. Spatial determinants, *e.g.* both the distribution of floral resources in the landscape and the way in which the pollinators interact with them, could therefore have a big impact on pollinator behaviour [36–40]. These are all determinants of how exactly a florally-transmitted pathogen is able to spread through the environment.

Unravelling how the spatial arrangement of the environment and how pollinators move through this environment and use flowers is therefore essential for understanding how pollinator health may be impacted [5]. In this paper, we present a spatially-explicit model of pollinator behaviour, considering how centrally-placed nests, each with multiple foraging bees, may interact with each other *via* an environment of transiently-present flowers, and consider the efficiency with which a hotspot-transmitted pathogen might propagate within that environment. Given that being able to recognise a potentially contaminated site is likely to be strongly selected for as a behavioural mechanism for avoiding pathogen transmission [5], we expand our model to consider a case where bees are able to deposit and detect scent marks on flowers before they expose themselves to contamination.

## RESULTS

### Infection model

Simulations suggest that the composition of foragers departing from the same nest location has little effect upon its likelihood of being infected, where changing the ratio of bees that either foraged close to home or went further away had no effect on infection speed (figure 1a). This lack of effect is probably because close-foraging bees have fewer flowers to visit within their smaller foraging radius when compared to far-foraging bees, and so are more likely to encounter an infection if it is present on at least one of the flowers within their range, while far-foraging bees are more likely to have an infected flower within the larger number of flowers within their foraging radius, but have a diluted risk of encountering them due to the increased number of flowers available to them. Similarly, the distance over which bees could see, once they had decided to stay in a square, had no effect (figure 1b). The distance that both far- and near-distance foragers were able to detect a new ‘target’ when seeking a new flower was influential however, and having more bees that could identify new ‘target’ flowers over longer distances meant that infections were more likely to spread quickly through the landscape (figures 1c and 1d).

**Figure 1.**
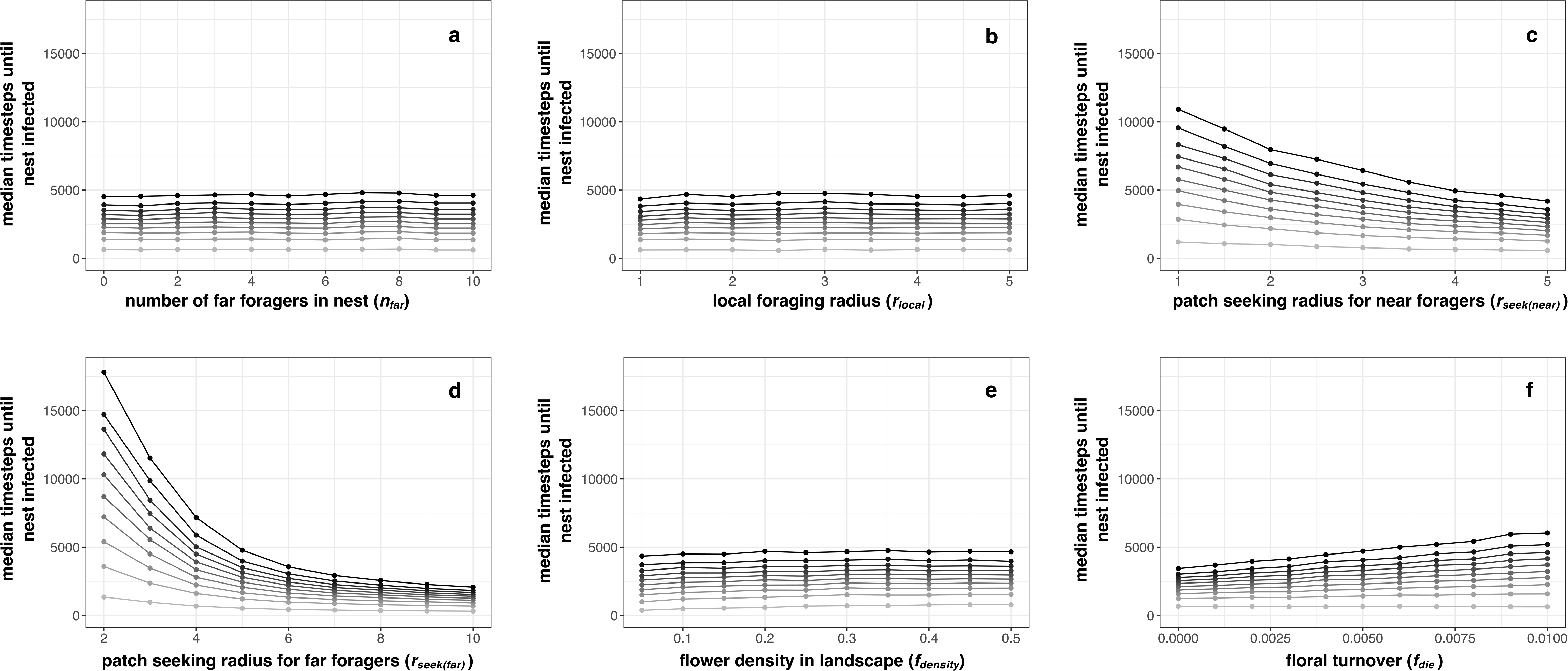
The spread of infection in response to model parameters. Median number of timesteps taken for at least one forager of a nest become infected *via* floral contact, when the following single parameters were systematically altered within a simulation: (a-d) behaviour of foragers, with a) number of far foragers in the nest; b) the local foraging radius; c) the flower-seeking radius of near foragers; d) the flower-seeking radius of far foragers; (e-f) floral parameters, e) the flower density in the landscape; and f) the rate of floral turnover. The colouring of the lines represents the order in which the ten nests present in a simulation were infected, with the lightest grey indicating the first nest to be infects, and the blackest line representing the last (tenth) nest to be infected, with the gradation of blackness indicating order.

Floral density appears to have a negligible impact on the spread of pathogens (figure 1e). Increasing the rate at which flowers died led to an increase in the time it took infections to spread, as with a faster floral turnover, tainted flowers were more likely to die, removing pathogen hotspots from the landscape(figure 1f).

The probability that a bee changed from foraging to returning home had an impact on infection, with bees that returned more quickly after leaving the nest leading to a great increase in the time taken until a nest had infected bees (figure 2a), presumably because the bees spend less time interacting with potentially contaminated flowers. Similarly, bees that were less likely to leave home led to pathogens taking longer to spread through the environment (figure 2b).

**Figure 2.**
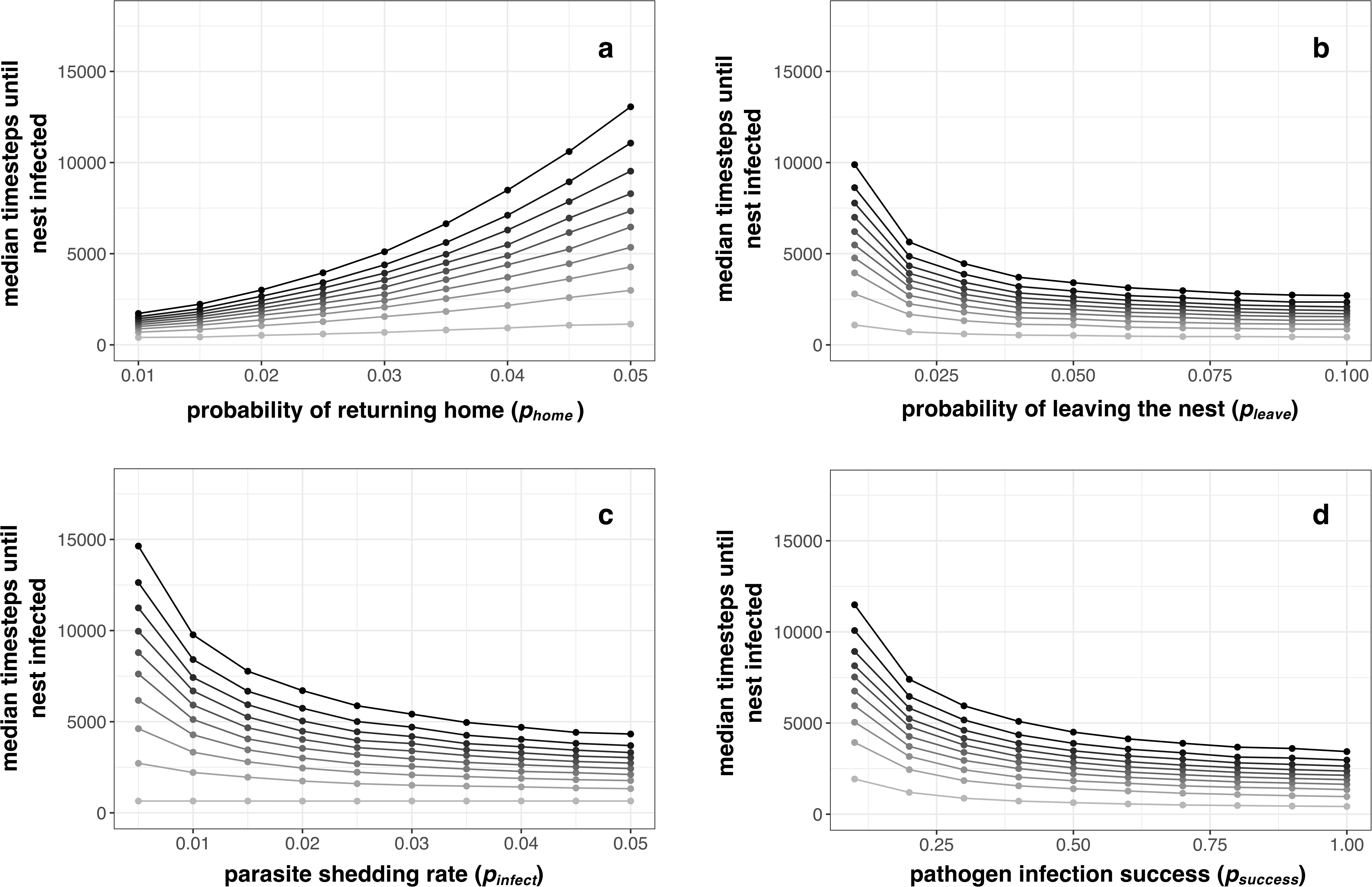
The spread of infection in response to model parameters that directly affect the spreading of the parasite. Median number of timesteps taken for at least one forager of a nest become infected *via* floral contact, when the following single parameters were systematically altered within a simulation: a) the probability that an individual within the nest decides to initiate flower visits and leaves the nest; b) the probability that an individual stops visiting flowers in order to return home; c) the rate of parasite shedding during a timestep when an infected individual is visiting a flower; and d) the probability that an individual is infected during a timestep when it is visiting an infected flower. The colouring of the lines represents the order in which the ten nests present in a simulation were infected, with the lightest grey indicating the first nest to be infects, and the blackest line representing the last (tenth) nest to be infected, with the gradation of blackness indicating order.

The rate at which infected bees shed pathogens generated the biggest effect on the spread of pathogens (figure 2c), where low shedding meant that the pathogen spread very slowly through the environment. Because of the shape of the curves shown, a pathogen that is shed rarely will benefit greatly from even a small increase in the rate at which it is shed, suggesting that the pathogens will be selected to increase shedding rate when they are rare. A similarly-shaped curve is seen with pathogen infection success (figure 2d), where pathogens that readily infect visitors tend to spread faster through the environment.

### Extended model, including scent mark deposition

Including scent marks generated little obvious difference in the model responses to many of the parameters considered, and results are not shown for these parameters for which there was no difference. However, scent mark deposition interacted with floral density (figure 3a), giving different results to the initial scent mark-free model where floral density had little effect on pathogen transmission dynamics (figure 1e). With scent marking occurring, the initial rate of transmission (shown in the paler low lines) tended to be faster in low-density environments, presumably because bees were more likely to come into contact with contaminated/tainted flowers that either had not yet been visited, or else had lost their scent marks. However, after the initial spread, infectious agents passed through the population more slowly in low-density environments, as contaminated/tainted flowers would be less likely to be investigated when the scent mark was still present, slowing down the spread of pathogens. For high-density environments, the reverse arguments could be made: more flowers present would mean that it took longer for bees to encounter contaminated/tainted flowers, but more flowers could also mean that bees would be less likely to encounter and avoid marked infected flowers when the infection was already common in the environment.

**Figure 3.**
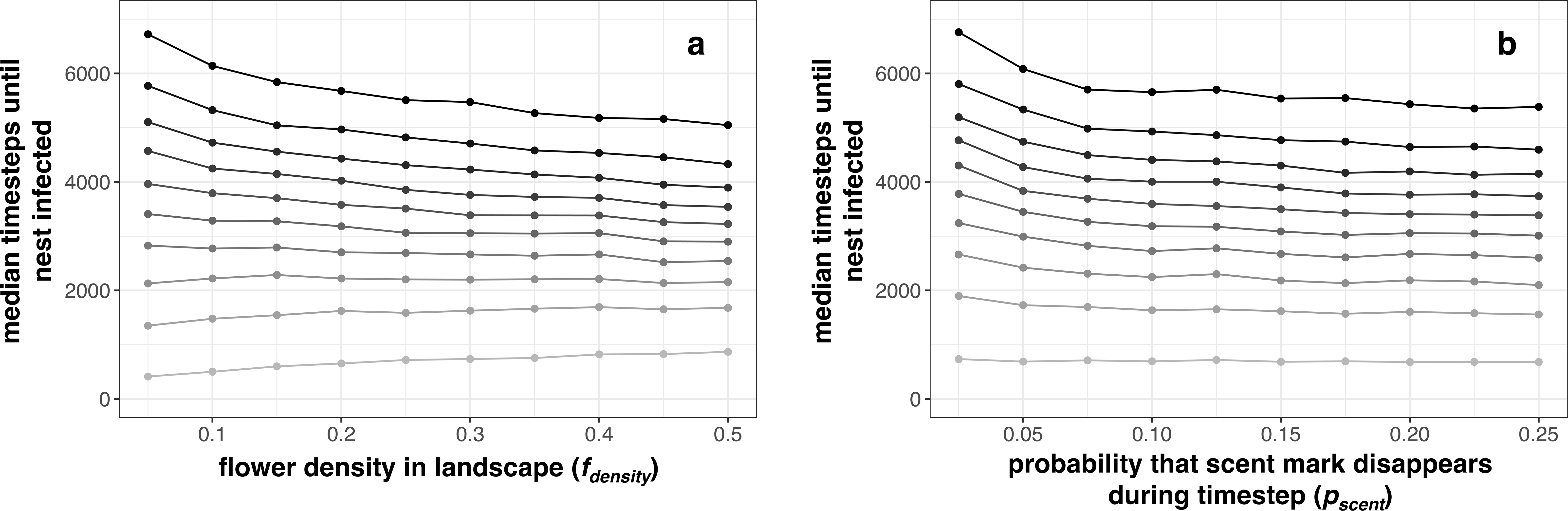
The spread of infection when bees deposited and responded to scent marks. Figures show median number of timesteps taken for nests to become infected, when single parameters were systematically altered within a simulation, considering a) flower density in the landscape, and b) the probability that a scent mark disappears during a timestep. The colouring of the lines represents the order in which the ten nests present in a simulation were infected, with the lightest grey indicating the first nest to be infects, and the blackest line representing the last (tenth) nest to be infected, with the gradation of blackness indicating order.

The longevity of the scent mark had little effect on the initial infection (figure 3b), which makes sense, given that the initial infection would have to come from an unmarked flower. Once the infection was present within the population, its rate of spread was reduced when scent marks were less likely to disappear.

## DISCUSSION

Here, we have presented a spatially-explicit model of pathogen transmission between pollinators *via* the flowers they visit. We found that the spatial arrangement of the environment and how pollinators move through this environment and use flowers has various impacts on pathogen transmission. Our results focus on the role that foraging behaviour has on the transmission of pathogens that spreads through flowers, a hitherto little quantified phenomenon. The biggest changes are associated with the likelihoods of leaving and staying in the nest and the distances travelled, and so it is likely that pollinators, most of which are constrained to return to a central nest, could be strongly impacted by the schedules they use for allocating foraging behaviour. Overall, the model we describe adds to our understanding of how pollinator movement in time and space contributes in variable ways to the spread of infections.

Our model considered the impact of some behavioural differences between foragers within a nest, by allowing some bees to travel further than others. Individual-based-models allow us to create structured populations of dissimilar individuals, which reflects the alloethism typically seen in bees [41]. Individual bee morphology may impact on likelihood of transmission [21], and body size has been shown to be important for both transmission and susceptibility in bumblebees [42]. We found that although differences in behaviour may be important, the composition of the nest may not have much impact on transmission, which suggests that alloethism within a nest is likely to reflect selective processes not related to pathogen-avoidance.

Our model did not allow individuals to change their behaviour over time, which may occur both in infected individuals [43] and in those encountering them ([44], but see [45]). Similarly, if a foraging bee was able to identify sites that potentially harbour pathogens (and therefore avoid them), this would impact on the spread of the pathogen (as for example seen in aversion behaviour in response to fungal spores [46] or the presence of *C. bombi* on a flower [47]): our model does not include this directly, but it would be assumed that being able to partially avoid pathogens would lead to their presence in the population and transmission rate being reduced.

Our model also demonstrated that scent marks impact on pathogen transmission as would be expected, but here we show that their effect may be dependent on both the longevity of the scent mark, the turnover of flowers in the environment, and the factors affecting pathogen viability and infectivity. Social foraging cues may override any avoidance cues about disease presence, as shown by [48,49], but it makes sense that scent marks should give indirect evidence that a possible foraging site may harbour an infectious agent.

Disease may affect the foraging behaviour and ecology of pollinators [50–52], and parasites can impose physical effects upon both cognition [53,54] and physical movement [55]. Changes in behaviour or movement could be accommodated within a spatially-explicit individual-based model, and careful parameterisation may be useful for exploring the impacts of different pollinator diseases. Floral traits may also be important for affecting transmission [56], and [21] showed experimentally that the number of reproductive structures present (but not the morphology or size) was related to transmission. Floral morphology may have an impact on the likelihood of pathogen deposition, for example bumblebees *B. impatiens* are more likely to defaecate on open plants rather than more complex plants [6]. Flower morphology can also affect exposure to the sun which impacts parasite survival [12], particularly as both floral behaviour [57,58] and petal surfaces [59–61] can influence the distribution of temperature gradients within the flower. Further work is needed to explore where particular pathogens are found within the flower, as their location (*e.g.* in the nectaries or on the petal surface) may impact on how the disease is transmitted [27]. It was unsurprising that increasing the rate of floral turnover in the model presented here made it harder for a pathogen to spread through a population, echoing empirical results [62]. Continuous trait modelling techniques [35] allow us to consider these specific criteria, and spatial individual-based models like the one presented in this paper could be adapted to explore handling and infectiveness differences between floral species, alongside exploration of differences caused by the species-specific biology of pathogens. Spatially explicit individual-based models would also allow us to consider non-biological environmental conditions such as temperature, which has been shown to affect the longevity of environmental *C. bombi* infective cells [63].

The density of pollinators within the environment may be important for increasing the transmission of a pathogen. For example, [64] shows high bumblebee host density can enhance on the transmission of viruses but not *C. bombi*. Our model does not explicitly consider nest density, but we can approach this by comparing models where foragers fly different distances from the nest (which has a big impact on transmission) and floral density (which has less impact). Different pollinator species may travel different mean distances within the environment [39,40,65,66], and this in turn is affected by the spatial distribution of floral resources relative to the nest, and so consideration of both pollinator density and species behavioural differences, as well as the density and spatial distribution of flowers may be important for understanding the dynamics of specific diseases. In the model, increasing floral density made it harder for infections to spread, presumably because foragers had more choice and were less likely to come into contact with an infected flower (and the limited lifespan of infected flowers would in turn have reduced contact still further). This echoes empirical findings in multi-species communities [67], where pathogen prevalence on flowers was lowest when floral abundance was highest.

Our results demonstrate that individual-based models can offer insights that may not easily be gained from standard compartmental models, given that pollinator-flower-pathogen interactions are likely to be spatially dependent upon pollinator behaviour, pollinator biology, and the spatial availability of flowers within the landscape. As well as spatial complexity, further landscape complexity could be introduced by considering interactions between multiple species – different co-occurring floral species may harbour different parasites [67], and the complexity of the interaction-network structure can drive the presence of pathogens [13]. Understanding these interactions could give us valuable insights into the dynamics and risks associated with pathogen spillover, which is known to occur [27]. This is particularly true when managed species are moved within the landscape, potentially carrying infections between areas [68]. Similarly, changes in the floral composition of the landscape may change the likelihood of different species interacting with each other: within a reward-poor landscape, cultivated wild flower meadows are pathogen-rich [69], and agricultural monocultures could act in a similar manner. Models incorporating these spatial, compositional and behavioural complexities are therefore important for understanding how plant-pollinator-pathogen interactions might be affected by both agricultural intensification and climate change [70].

## METHODS

### Overview of models

Here, we consider a spatially-explicit individual-based model, run in *Netlogo* 6.4.0 [71]. Model code, along with all the data and *R* code used in this paper, is freely available at [72]. The model considers how pathogens can spread through a landscape that has a random distribution of flowers within it, by assuming that a number of social bee nests are randomly scattered within this landscape. Each nest contains a number of individual bees (for simplicity, we assume ten foragers in each nest), who choose and then conduct their movements within the landscape based on a series of behavioural rules that are influenced by what they can experience of the environment (based in this model on being able to see and choose between flowers within a maximum radius from their current position), alongside their internal state (in this model, dictated by a probabilistic preference for returning home and a probabilistic preference for switching between flower-seeking behaviour and foraging behaviour). Individual bees within a nest can have differing sensory abilities (that influence the distance over which they can detect flowers), echoing the differences in foraging behaviour seen within real-life nests of some social bees, and differences in body size of sympatric solitary and social bee species [73–75]. In this model, bees that come into contact with flowers contaminated with a pathogen (here, the flowers are labelled as ‘tainted’ to avoid ambiguity in terminology) have a chance of becoming infected that is directly related to the time they spend on the flower, and then stochastically shed the infection at a fixed probability when they visit subsequent flowers.

We simulate an infection event by seeding the environment with a single infection, and then track how the disease spreads through the bee population by tracking the time at which it spreads to new nests, assuming that the infection process is limited solely to transfer of the pathogen when the bee is on a flower, with no transmission occurring within the nest. We also consider the effects of a changeable environment by allowing flowers to die and bloom within the environment at each timestep, meaning that contaminated flowers can be removed (although we assume that there is always at least one site of pathogen transmission present in the environment, as this study is not concerned with population processes, but rather how infections are able to spread). Consequently, we assume that the timescale of our simulations is short enough to ensure that individual bees do not die, and so we do not include a birth/death process for the bees.

In a second set of models, all foraging bees also deposit a scent mark on the flowers that they visit [33,34]. Subsequent foragers are able to detect scent marks when they are in immediate proximity of the flower, and will alter their behaviour to avoid landing on the flower (and therefore missing infection, if there was an infection present on the flower). A scent mark is assumed to decay, and flowers that are contaminated and have persisted in the environment are likely to lose their marks and appear ‘clean’ to a visitor.

### Environment

The model is set within a 101 × 101 square grid over the surface of a torus, such that the edges of the grid are attached, and the edges of each square are assumed to be 1 distance unit in length. At any moment in time, each square can contain either a single flower, or no flower. The initial flower density *f_density_*defines the proportion of the landscape that has flowers: at the beginning of a simulation, each square has a probability *f_density_* of having a single untainted flower in it (noting that we subsequently refer to this probability as ‘flower density’), otherwise the square is recorded as being empty. Once these are placed, then *n_nest_*nests are placed randomly on different squares in the grid (and if there is a flower already present in a nest square, it is deleted). Each nest is populated with *n_bees_* bees, who are initially orientated in randomly chosen directions and are all initially labelled as being uninfected. Each bee is labelled with the location of its nest, so it knows which square it needs to return to. Finally, one flower is randomly chosen to become the sole source of infection within the environment, and the square containing it is labelled as being tainted with the pathogen.

### Flower turnover

Within a simulation, we consider each flower to have a fixed probability *f_die_* of dying during each timestep (this is done before the bees start making decisions and moving). This probability does not consider whether a flower is tainted or not (with the exception of where the tainted flower is the only source of infection in the environment, such as at the start of the simulation – if this is the case, the solitary tainted flower is assumed to persist, and we only allow it a probabilistic ability to die once the infection has spread to other flowers). If a flower dies, it is immediately removed from the landscape, and the square it occupied is recorded as being empty. We note the number of flowers that die within a timestep, denoted *n_die_*. Within a timestep, once all decisions have been made about whether flowers have died or persisted, we then allow *n_die_* untainted flowers to bloom, scattering them randomly across all of the empty squares (*e.g.* those containing neither a flower nor a nest) – this means that *n_die_* remains constant throughout a simulation.

### Bee movement

At the start of each timestep, bees can be in one of four behavioural states (‘nesting’, ‘seeking flowers’, ‘local foraging’, or ‘returning home’), dependent upon both its current location and its previous action. During its turn, the bee will firstly make a decision about whether to change or remain in its initial behavioural state, and may then make a movement dependent upon this updated state. We allow bees to revisit flowers as happen in their natural behaviour. Bees conduct multiple foraging bouts during a simulation, and can leave and return to the nest an unlimited number of times. Because our simulations involve bees being able to identify flowers in their environment that may take multiple timesteps (and therefore decision-points) to reach, we use the term ‘target’ to refer to a site that has been identified and that behaviour is being directed towards (and so bees may or may not have a ‘target’ set at the start of each timestep).

If the bee is currently on the nest square, its behavioural state at the start of the timestep is ‘nesting’. It will update its behavioural state to ‘seeking flowers’ with probability *p_leave_*, at the same time choosing a new random heading, and it is assumed here that the bee will leave the nest without information about the environment, and so a target is not set at this moment (so we implicitly make the assumption that information is not shared about flower location, which would be typical in non-honeybee species). Otherwise, the bee remains in the nest, and its behavioural state remains as ‘nesting’.

If the bee is not on the nest square and its behavioural state at the start of the timestep is ‘seeking flowers’, it first decides whether it is time to return to the nest with probability *p_return_*, where the heading of the bee is shifted to the quickest route back to its nest and its behavioural state is updated to ‘returning home’. If the bee has not decided to return home, it then checks whether it has a target set. If it does not have a target, it randomly chooses an unoccupied flower visible within a radius *r_seek_*, sets this location as its target, and orientates itself to be able to move to this target by the shortest route (or, if no unoccupied flower is visible within radius *r_seek_*, it chooses a random heading without setting a target). If instead the bee already has a target set at the start of its turn, it checks whether it has reached that target. If it has, it updates its behavioural state to ‘local foraging’, and orientates itself randomly with no target (meaning that it then needs to spend a turn making local movement decisions once it has reached the site in which it means to forage).

If the bee is not on the nest square and its behavioural state at the start of the timestep is ‘local foraging’, it first decides whether it is time to return to the nest with probability *p_return_*, where the heading of the bee is shifted to the quickest route back to its nest and its behavioural state is updated to ‘returning home’. It then uses a density-dependent rule to decide whether it switches back to ‘seeking flowers’ with no target set and a random heading, where the switch occurs with probability

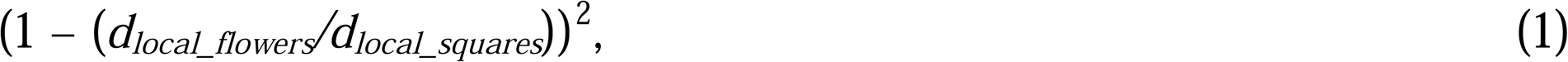

where *d_local_flowers_* is the number of unoccupied flowers visible within radius *r_local_*of the bee, and *d_local_squares_* is the number of complete or partially complete squares (discounting the current square) visible within radius *r_local_* of the bee. If the bee chooses to remain as local foraging, if it does not have a target set it will choose an unoccupied square containing a flower within radius *r_local_*, record this as its target, and orientate towards it (or, if no unoccupied flower is visible within radius *r_seek_*, it chooses a random heading without setting a target). If it already has a target set, it will check whether it currently occupies the target; if it does, it will choose an unoccupied square containing a flower within radius *r_local_*, record this as its target, and orientate towards it (or, if no unoccupied flower is visible within radius *r_seek_*, it chooses a random heading without setting a target). If instead it already has a target set, but does not currently occupy that square, it will orientate itself towards the target.

Finally, if the bee is not on the nest square and its behavioural state at the start of the timestep is ‘returning home’, it will first check whether it currently occupies the nest square. If it does, it updates its behavioural state to ‘nesting’, otherwise, it orientates itself to take the shortest route back to the nest.

Once the bee has decided on its action, it will then either move forwards one distance unit (if its behavioural state is ‘seeking flowers’, ‘local foraging’ or ‘returning home’), or remain in place (if its behavioural state is ‘nesting’). Because movement decisions are made before movement happens, we note that there is a very small chance that two bees arrive on the same flower at the same time, which could occur if both are one distance unit away from the flower at the start of a timestep and both decide to move to the flower.

### Different bee types in a nest

A nest can consist of a heterogeneous mix of different individuals, who forage different distances away. For simplicity, we consider nests as being able to consist of two types of individuals: *n_near_* near-distance foragers, who have a low personal *r_seek_* value, and *n_far_* far-distance foragers, who have a higher personal *r_seek_* value.

### Infection

If an uninfected bee is in a square containing a contaminated flower, it is assumed to become infected with probability *p_success_*, which is the probability that the pathogen is successfully passed on to the visiting bee during a timestep. If the bee is already infected, but the flower is not tainted, the flower becomes tainted with the pathogen with probability *p_infect_*, or otherwise remains untainted. Within the simulation, actions on both the bee and flower occur at the start of a movement decision, before any movement is made. Consequently, in the very small chance of two bees arriving at the flower at the same time, transmission from a bee carrying the pathogen to one without could only occur with the multiple timesteps necessary to deposit and then pick up the pathogen.

### Time step

During a simulation timestep, all flower deaths and replacements are conducted before all other actions. Following this, the order in which the bees move is randomised, such that all bees make one behaviour decision (and possible movement) during a timestep. Each selected bee in turn decides which behaviour to conduct and makes a movement as required (with any resulting changes to the environment such as infecting a flower also being conducted), before any action is taken by the next bee.

### Simulations and model exploration

For all simulations, we assumed *n_nest_* = 10 nests are randomly distributed through the environment, each containing *n_bees_* = 10 bees. To investigate the model, each parameter was systematically varied in turn. 5000 different environments were generated for each parameter being investigated, where the landscape and non-investigated parameters were randomly generated from the following distributions (assuming the parameter being systematically investigated was missed from these): *n_far_* was varied between 0 and 10, with *n_near_* = 10 – *n_far_*; *r_local_*took an integer value between 3 and 6; *r_seek_* took an integer value between 3 and 6 for near foragers (denoted *r_seek(near)_*), and (assuming *r_seek(near, simulation)_* has already been generated for the simulation) an integer value between *r_seek(near,simulation)_*and *r_seek(near,simulation)_* + 4 for far foragers (denoted *r_seek(far)_*_)_; the initial flower density *f_density_* varied between 0.05 and 0.50, and the flower turnover *f_die_* varied between 0 and 0.01; the probability *_pinfect_* that an infected bee shed the pathogen when it visited a flower varied between 0.005 and 0.01, and the pathogen infection success *p_success_* varied between 0.1 and 1; bees chose to return home with a value of *p_home_*between 0.01 and 0.05, and chose to leave the nest with a value of *p_leave_*between 0.01 and 0.05. When choosing values between the ranges given, we assumed uniform distributions, so any value in the range was equally likely to be chosen. The single parameters being systematically investigated took the following values: *n_far_*= (0, 1, … 10); *r_local_* = (1, 1.5, …, 5); *r_seek(near)_*= (1, 1.5, …, 5) for near-distance foragers, or *r_seek(far)_*+ (2, 3, …, 10) for far-distance foragers (with the added limitation that within each simulation, *r_seek(near)_* was then sampled from a uniform distribution with range (1, *r_seek(far)_*); *f_density_*= (0.05, 0.075, …, 0.5); *f_die_* = (0, 0.001, …, 0.01); *p_infect_* = (0.005, 0.01, …, 0.05); *p_success_* = (0.1, 0.2, …, 1); *p_home_* = (0.01, 0.015, …, 0.05); and *p_leave_* = (0.01, 0.02, …, 0.1).

In order to quantify the spread of pathogens through the landscape, we recorded the timestep at which a nest is first visited by an infected resident, according to how many nests had already been infected. We assume that the simulation starts at timestep 1. Each simulation was ended either as soon as the final nest had infected individuals, or if 100,000 timesteps had been simulated without all nests containing infected bees. If the maximum length of the simulation was reached without full infection, the nests that did not have infected bees were recorded as being pathogen-free, and were not included in the analysis. This means that the statistics calculated are likely to be smaller than the true values, but given that very few nests were uninfected at the ends of most simulations, this will have little effect on the effects described.

For each of the focus parameter values within the simulations, we calculated the median values of the timesteps at which the first, second, third *etc*. nests housed infected bees. The means were also calculated (not shown), which showed similar trends, but with much higher values due to some of the simulations taking extremely long times to finish – caused either by randomly generated parameter sets that meant either that the bees were unable to explore the landscape well, or an initial infection was placed at a point where it could not be discovered by the bees). We also explored spatially-smaller (51 × 51) versions of the model, including a version where the density-dependent probability shown in eqn. 1 was replaced with (1 – (*d_local_flowers_/d_local_squares_*)), and the results (not shown) were qualitatively similar to those described. All data manipulation was conducted within *R* 4.3.0 [76], using *ggplot2* 3.4.4 [77] for visualisation.

### Scent mark deposition

An additional full set of simulations were conducted where the assumptions made above were followed, but where bees were also able to deposit and to detect previously-deposited scent marks on floral surfaces. At the start of the simulation, it was assumed that no flowers had already been visited by bees, and when new flowers were created, we also assumed they had not already been visited. All these flowers therefore did not initially have scent marks. Scent marks were only present on flowers that had been visited by bees, as described below.

At the beginning of time step, after the floral birth/death process had been conducted but before the bees present made their behavioural decisions and moves, we updated the scent marks on the flowers. If an individual flower had possessed a scent mark in the previous step, the scent mark disappeared with probability *p_scent_*, giving an exponential decay to the chemical mark present. If a scent mark disappeared in a given time step, it was assumed to have gone from that point onwards, with flowers only able to regain scent marks when they were subsequently visited by a bee. It is worth emphasising that scent marks and parasite infections are not completely correlated – the initial source of infection in the model will not have a scent mark present, while a scent mark does not necessarily indicate the presence of a pathogen, and pathogens may be found on ‘clean’ flowers that have lost their scent marks.

The individual behavioural rules were similar to those described above. For bees conducting ‘flower seeking’ or ‘local foraging’ behaviours, the bees went through their normal decision processes as described above. Having done this, but before moving, the bees monitored the square one distance unit ahead of them; if no scent mark was present, they moved one step forwards as described. If however the square ahead contained a flower (which would have been set as the target flower within these behaviours) and the flower had a scent mark deposited on it, the bee then redefined the flower as not being its target, and moved one step forward (simulating visiting the square with the flower in it, but not interacting with the flower having identified it as being one that has been recently visited). This meant that the bee started the next period as not having a target set, which would allow it to rechoose and reorientate itself within the environment.

When a bee reached a flower it identified as its target (meaning that the flower did not carry a scent mark already), the flower had a scent mark deposited on it.

A suite of simulations were conducted similar to those described for the previous model, with *p_scent_* taking a random value between 0.025 and 0.250 in each simulation set. An extra set of simulations explored the scent mark decay by randomly altering the parameter set as described for the previous model, with each parameter set being run for *p_scent_* = (0.025, 0.050, …, 0.250). All other model assumptions and data harvested were as described for the previous model.

## FUNDING

E.N. acknowledges support from UK Research and Innovation (Future Leaders Fellowship awarded to E.N., MR/T021691/1). C.B. acknowledges support from the Spanish Ministry of Science and Innovation (‘Ramón y Cajal’ contract awarded to C.B., RYC2020-028962-I) and from the Spanish Research Agency (project ref.: PID2022-143255OA-I00). N.H.dI. acknowledges support from the Biotechnology and Biological Sciences Research Agency (BB/N000625/1).

## Notes

https://doi.org/10.5281/zenodo.15005303

### Competing Interest Statement

The authors have declared no competing interest.

## REFERENCES

1. Kotwal G, Cannon JL. Environmental persistence and transfer of enteric viruses. Curr Opin Virol. 2014;4: 37–43. doi:10.1016/j.coviro.2013.12.003

2. La Rosa G, Fratini M, Libera SD, Iaconelli M, Muscillo M. Viral infections acquired indoors through airborne, droplet or contact transmission. Ann Ist Super Sanità. 2013;49: 124–132. doi:10.4415/ANN_13_02_03

3. Fenton A, Rands SA. Optimal parasite foraging strategies: a state-dependent approach. Int J Parasitol. 2004;34: 813–821. doi:10.1016/j.ijpara.2004.02.003

4. Durrer S, Schmid-Hempel P. Shared use of flowers leads to horizontal pathogen transmission. Proc R Soc B. 1994;258: 299–302. doi:10.1098/rspb.1994.0176

5. Nicholls E, Rands SA, Botías C, Hempel de Ibarra N. Flower sharing and pollinator health: a behavioural perspective. Phil Trans R Soc B. 2022;377: 20210157. doi:10.1098/rstb.2021.0157

6. Bodden JM, Hazlehurst JA, Wilson Rankin EE. Floral traits predict frequency of defecation on flowers by foraging bumble bees. J Insect Sci. 2019;19(5): 2. doi:10.1093/jisesa/iez091

7. Davis AE, Deutsch KR, Torres AM, Mata Loya MJ, Cody LV, Harte E, et al. *Eristalis* flower flies can be mechanical vectors of the common trypanosome bee parasite, *Crithidia bombi*. Sci Rep. 2021;11: 15852. doi:10.1038/s41598-021-95323-w

8. Adler LS, Barber NA, Biller OM, Irwin RE. Flowering plant composition shapes pathogen infection intensity and reproduction in bumble bee colonies. Proc Natl Acad Sci USA. 2020;117: 11559–11565. doi:10.1073/pnas.2000074117

9. McArt SH, Koch H, Irwin RE, Adler LS. Arranging the bouquet of disease: floral traits and the transmission of plant and animal pathogens. Ecol Lett. 2014;17: 624–636. doi:10.1111/ele.12257

10. Pinilla-Gallego MS, Ng WH, Amaral VE, Irwin RE. Floral shape predicts bee–parasite transmission potential. Ecology. 2022;103: e3730. doi:10.1002/ecy.3730

11. Van Wyk JI, Lynch A-M, Adler LS. Manipulation of multiple floral traits demonstrates role in pollinator disease transmission. Ecology. 2023;104: e3866. doi:10.1002/ecy.3866

12. Figueroa LL, Blinder M, Grincavitch C, Jelinek A, Mann EK, Merva LA, et al. Bee pathogen transmission dynamics: deposition, persistence and acquisition on flowers. Proc R Soc B. 2019;286: 20190603. doi:10.1098/rspb.2019.0603

13. Figueroa LL, Grab H, Ng WH, Myers CR, Graystock P, McFrederick QS, et al. Landscape simplification shapes pathogen prevalence in plant-pollinator networks. Ecol Lett. 2020;23: 1212–1222. doi:10.1111/ele.13521

14. Ellner SP, Ng WH, Myers CR. Individual specialization and multihost epidemics: disease spread in plant-pollinator networks. Am Nat. 2020;195: E118–E131. doi:10.1086/708272

15. Heinrich B. Do bumblebees forage optimally, and does it matter? Am Zool. 1983;23: 273–281. doi:10.1093/icb/23.2.273

16. Goulson D. Why do pollinators visit proportionally fewer flowers in large patches? Oikos. 2000;91: 485–492. doi:10.1034/j.1600-0706.2000.910309.x

17. Baird E, Tichit P, Guiraud M. The neuroecology of bee flight behaviours. Curr Opin Insect Sci. 2020;42: 8–13. doi:10.1016/j.cois.2020.07.005

18. Greggers U, Menzel R. Memory dynamics and foraging strategies of honeybees. Behav Ecol Sociobiol. 1993;32: 17–29. doi:10.1007/BF00172219

19. Lihoreau M, Raine NE, Reynolds AM, Stelzer RJ, Lim KS, Smith AD, et al. Radar tracking and motion-sensitive cameras on flowers reveal the development of pollinator multi-destination routes over large spatial scales. PLoS Biol. 2012;10: e1001392. doi:10.1371/journal.pbio.1001392

20. Rands SA, Whitney HM, Hempel de Ibarra N. Multimodal floral recognition by bumblebees. Curr Opin Insect Sci. 2023;59: 101086. doi:10.1016/j.cois.2023.101086

21. Adler LS, Michaud KM, Ellner SP, McArt SH, Stevenson PC, Irwin RE. Disease where you dine: plant species and floral traits associated with pathogen transmission in bumble bees. Ecology. 2018;99: 2535–2545. doi:10.1002/ecy.2503

22. Cisarovsky G, Schmid-Hempel P. Combining laboratory and field approaches to investigate the importance of flower nectar in the horizontal transmission of a bumblebee parasite. Entomol Exp Appl. 2014;152: 209–215. doi:10.1111/eea.12218

23. Graystock P, Goulson D, Hughes WOH. Parasites in bloom: flowers aid dispersal and transmission of pollinator parasites within and between bee species. Proc R Soc B. 2015;282: 20151371. doi:10.1098/rspb.2015.1371

24. Schwarz HH, Huck K. Phoretic mites use flowers to transfer between foraging bumblebees. Insectes Soc. 1997;44: 303–310. doi:10.1007/s000400050051

25. Peck DT, Smith ML, Seeley TD. *Varroa destructor* mites can nimbly climb from flowers onto foraging honey bees. PLoS One. 2016;11: e0167798. doi:10.1371/journal.pone.0167798

26. Alger SA, Burnham PA, Brody AK. Flowers as viral hot spots: Honey bees (*Apis mellifera*) unevenly deposit viruses across plant species. PLoS One. 2019;14: e0221800. doi:10.1371/journal.pone.0221800

27. Alger SA, Burnham PA, Boncristiani HF, Brody AK. RNA virus spillover from managed honeybees (*Apis mellifera*) to wild bumblebees (*Bombus* spp.). PLoS One. 2019;14: e0217822. doi:10.1371/journal.pone.0217822

28. Burnham PA, Alger SA, Case B, Boncristiani H, Hébert-Dufresne L, Brody AK. Flowers as dirty doorknobs: Deformed wing virus transmitted between *Apis mellifera* and *Bombus impatiens* through shared flowers. J Appl Ecol. 2021;58: 2065–2074. doi:10.1111/1365-2664.13962

29. Singh R, Levitt AL, Rajotte EG, Holmes EC, Ostiguy N, vanEngelsdorp D, et al. RNA viruses in Hymenopteran pollinators: evidence of inter-taxa virus transmission *via* pollen and potential impact on non-*Apis* Hymenopteran species. PLoS One. 2010;5: e14357. doi:10.1371/journal.pone.0014357

30. Evison SE, Jensen AB. The biology and prevalence of fungal diseases in managed and wild bees. Curr Opin Insect Sci. 2018;26: 105–113. doi:10.1016/j.cois.2018.02.010

31. Takeda K, Sakai S. Extended benefits of pollinator-mediated microbial dispersal among flowers. Ecol Res. 2022;37: 481–484. doi:10.1111/1440-1703.12326

32. Eigenbrode SD, Bosque-Pérez NA, Davis TS. Insect-borne plant pathogens and their vectors: ecology, evolution, and complex interactions. Annu Rev Entomol. 2018;63: 169–191. doi:10.1146/annurev-ento-020117-043119

33. Pearce RF, Giuggioli L, Rands SA. Bumblebees can discriminate between scent-marks deposited by conspecifics. Sci Rep. 2017;7: 43872. doi:10.1038/srep43872

34. Stout JC, Goulson D. The use of conspecific and interspecific scent marks by foraging bumblebees and honeybees. Anim Behav. 2001;62: 183–189. doi:10.1006/anbe.2001.1729

35. Truitt LL, McArt SH, Vaughn AH, Ellner SP. Trait-based modeling of multihost pathogen transmission: plant-pollinator networks. Am Nat. 2019;193: E149–E167. doi:10.1086/702959

36. Becher MA, Grimm V, Thorbek P, Horn J, Kennedy PJ, Osborne JL. BEEHAVE: a systems model of honeybee colony dynamics and foraging to explore multifactorial causes of colony failure. J Appl Ecol. 2014;51: 470–482. doi:10.1111/1365-2664.12222

37. Becher MA, Twiston-Davies G, Penny TD, Goulson D, Rotheray EL, Osborne JL. Bumble-BEEHAVE: A systems model for exploring multifactorial causes of bumblebee decline at individual, colony, population and community level. J Appl Ecol. 2018;55: 2790–2801. doi:10.1111/1365-2664.13165

38. Rands SA. Landscape fragmentation and pollinator movement within agricultural environments: a modelling framework for exploring foraging and movement ecology. PeerJ. 2014;2: e269. doi:10.7717/peerj.269

39. Rands SA, Whitney HM. Effects of pollinator density-dependent preferences on field margin pollination in the midst of agricultural monocultures: a modelling approach. Ecol Model. 2010;221: 1310–1316. doi:10.1016/j.ecolmodel.2010.01.014

40. Rands SA, Whitney HM. Field margins, foraging distances and their impacts on nesting pollinator success. PLoS One. 2011;6: e25971. doi:10.1371/journal.pone.0025971

41. Goulson D, Peat J, Stout JC, Tucker J, Darvill B, Derwent LC, et al. Can alloethism in workers of the bumblebee, *Bombus terrestris*, be explained in terms of foraging efficiency? Anim Behav. 2002;64: 123–130. doi:10.1006/anbe.2002.3041

42. Van Wyk JI, Amponsah ER, Ng WH, Adler LS. Big bees spread disease: body size mediates transmission of a bumble bee pathogen. Ecology. 2021;102: e03429. doi:10.1002/ecy.3429

43. Goblirsch M, Huang ZY, Spivak M. Physiological and behavioral changes in honey bees (*Apis mellifera*) induced by *Nosema ceranae* infection. PLoS One. 2013;8: e58165. doi:10.1371/journal.pone.0058165

44. Baracchi D, Fadda A, Turillazzi S. Evidence for antiseptic behaviour towards sick adult bees in honey bee colonies. J Insect Physiol. 2012;58: 1589–1596. doi:10.1016/j.jinsphys.2012.09.014

45. Murray ZL, Keyzers RA, Barbieri RF, Digby AP, Lester PJ. Two pathogens change cuticular hydrocarbon profiles but neither elicit a social behavioural change in infected honey bees, *Apis mellifera* (Apidae: Hymenoptera). Austral Entomol. 2016;55: 147–153. doi:10.1111/aen.12165

46. Yousefi B, Fouks B. The presence of a larval honey bee parasite, *Ascosphaera apis*, on flowers reduces pollinator visitation to several plant species. Acta Oecol. 2019;96: 49–55. doi:10.1016/j.actao.2019.03.006

47. Fouks B, Lattorff HMG. Recognition and avoidance of contaminated flowers by foraging bumblebees (*Bombus terrestris*). PLoS One. 2011;6: e26328. doi:10.1371/journal.pone.0026328

48. Fouks B, Lattorff HMG. Social scent marks do not improve avoidance of parasites in foraging bumblebees. J Exp Biol. 2013;216: 285–291. doi:10.1242/jeb.075374

49. Fouks B, Robb EG, Lattorff HMG. Role of conspecifics and personal experience on behavioral avoidance of contaminated flowers by bumblebees. Curr Zool. 2019;65: 447–455. doi:10.1093/cz/zoy099

50. Francis JS, Tatarko AR, Richman SK, Vaudo AD, Leonard AS. Microbes and pollinator behavior in the floral marketplace. Curr Opin Insect Sci. 2021;44: 16–22. doi:10.1016/j.cois.2020.10.003

51. Koch H, Brown MJ, Stevenson PC. The role of disease in bee foraging ecology. Curr Opin Insect Sci. 2017;21: 60–67. doi:10.1016/j.cois.2017.05.008

52. Schmid-Hempel P, Stauffer H-P. Parasites and flower choice of bumblebees. Anim Behav. 1998;55: 819–825. doi:10.1006/anbe.1997.0661

53. Gómez-Moracho T, Heeb P, Lihoreau M. Effects of parasites and pathogens on bee cognition. Ecol Entomol. 2017;42(S1): 51–64. doi:10.1111/een.12434

54. Gómez-Moracho T, Durand T, Lihoreau M. The gut parasite *Nosema ceranae* impairs olfactory learning in bumblebees. J Exp Biol. 2022;225: jeb244340. doi:10.1242/jeb.244340

55. Wells T, Wolf S, Nicholls E, Groll H, Lim KS, Clark SJ, et al. Flight performance of actively foraging honey bees is reduced by a common pathogen. Environ Microbiol Rep. 2016;8: 728–737. doi:10.1111/1758-2229.12434

56. Fouks B, Wagoner KM. Pollinator parasites and the evolution of floral traits. Ecol Evol. 2019;9: 6722–6737. doi:10.1002/ece3.4989

57. Kevan PG. Sun-tracking solar furnaces in high arctic flowers: significance for pollination and insects. Science. 1975;189: 723–726. doi:10.1126/science.189.4204.723

58. Vandenbrink JP, Brown EA, Harmer SL, Blackman BK. Turning heads: the biology of solar tracking in sunflower. Plant Sci. 2014;224: 20–26. doi:10.1016/j.plantsci.2014.04.006

59. Harrap MJM, Rands SA, Hempel de Ibarra N, Whitney HM. The diversity of floral temperature patterns, and their use by pollinators. eLife. 2017;6: e31262. doi:10.7554/eLife.31262

60. Harrap MJM, de Vere N, Hempel de Ibarra N, Whitney HM, Rands SA. Variations of floral temperature in changing weather conditions. Ecol Evol. 2024;14: e11651. doi:10.1002/ece3.11651

61. Harrap MJM, Hempel de Ibarra N, Whitney HM, Rands SA. Floral temperature patterns can function as floral guides. Arthropod-Plant Interact. 2020;14: 193–206. doi:10.1007/s11829-020-09742-z

62. Piot N, Eeraerts M, Pisman M, Claus G, Meeus I, Smagghe G. More is less: mass-flowering fruit tree crops dilute parasite transmission between bees. Int J Parasitol. 2021;51: 777–785. doi:10.1016/j.ijpara.2021.02.002

63. Wolmuth-Gordon HS, Brown MJF. Transmission of a bumblebee parasite is robust despite parasite exposure to extreme temperatures. Ecol Evol. 2023;13: e10379. doi:10.1002/ece3.10379

64. Bailes EJ, Bagi J, Coltman J, Fountain MT, Wilfert L, Brown MJF. Host density drives viral, but not trypanosome, transmission in a key pollinator. Proc R Soc B. 2020;287: 20191969. doi:10.1098/rspb.2019.1969

65. Greenleaf SS, Williams NM, Winfree R, Kremen C. Bee foraging ranges and their relationship to body size. Oecologia. 2007;153: 589–596. doi:10.1007/s00442-007-0752-9

66. Knight ME, Osborne JL, Sanderson RA, Hale RJ, Martin AP, Goulson D. Bumblebee nest density and the scale of available forage in arable landscapes. Insect Conserv Divers. 2009;2: 116–124.

67. Graystock P, Ng WH, Parks K, Tripodi AD, Muñiz PA, Fersch AA, et al. Dominant bee species and floral abundance drive parasite temporal dynamics in plant-pollinator communities. Nat Ecol Evol. 2020;4: 1358–1367. doi:10.1038/s41559-020-1247-x

68. Manley R, Boots M, Wilfert L. Emerging viral disease risk to pollinating insects: ecological, evolutionary and anthropogenic factors. J Appl Ecol. 2015;52: 331–340. doi:10.1111/1365-2664.12385

69. Piot N, Meeus I, Kleijn D, Scheper J, Linders T, Smagghe G. Establishment of wildflower fields in poor quality landscapes enhances micro-parasite prevalence in wild bumble bees. Oecologia. 2019;189: 149–158. doi:10.1007/s00442-018-4296-y

70. Proesmans W, Albrecht M, Gajda A, Neumann P, Paxton RJ, Pioz M, et al. Pathways for novel epidemiology: plant–pollinator–pathogen networks and global change. Trends Ecol Evol. 2021;36: 623–636. doi:10.1016/j.tree.2021.03.006

71. Wilensky U. NetLogo [online: http://ccl.northwestern.edu/netlogo/]. Evanston: Center for Connected Learning and Computer-based Modeling, Northwestern University; 1999.

72. Rands SA, Botías C, Nicholls E, Hempel de Ibarra N. A spatially explicit model of pollinator-plant-pathogen interactions: Netlogo code and datasets. Zenodo. 2025; 10.5281/zenodo.15005303. doi:10.5281/zenodo.15005303

73. Hagbery J, Nieh JC. Individual lifetime pollen and nectar foraging preferences in bumble bees. Naturwissenschaften. 2012;99: 821–832. doi:10.1007/s00114-012-0964-7

74. Spaethe J, Weidenmüller A. Size variation and foraging rate in bumblebees (*Bombus terrestris*). Insectes Soc. 2002;49: 142–146. doi:10.1007/s00040-002-8293-z

75. Roulston TH, Cane JH. The effect of diet breadth and nesting ecology on body size variation in bees (Apiformes). J Kans Entomol Soc. 2000;73: 129–142.

76. R Development Core Team. R: a language and environment for statistical computing. Vienna: R Foundation for Statistical Computing; 2024.

77. Wickham H. ggplot2: elegant graphics for data analysis. New York: Springer-Verlag; 2009.

